# The ATP-dependent chromatin remodelling enzyme Uls1 prevents Topoisomerase II poisoning

**DOI:** 10.1101/412783

**Authors:** Amy Swanston, Katerina Zabrady, Helder C. Ferreira

## Abstract

Topoisomerase II (Top2) is an essential enzyme that decatenates DNA via a transient Top2-DNA covalent intermediate. This intermediate can be stabilised by a class of drugs termed Top2 poisons, resulting in massive DNA damage. Thus, Top2 activity is a double-edged sword that needs to be carefully controlled to maintain genome stability. We show that Uls1, an ATP-dependent chromatin remodelling (Snf2) enzyme, can alter Top2 chromatin binding and prevent Top2 poisoning in yeast. Deletion mutants of *ULS1* are hypersensitive to the Top2 poison acriflavine (ACF), activating the DNA damage checkpoint. We map Uls1’s Top2 interaction domain and show that this, together with its ATPase activity, is essential for Uls1 function. By performing ChIP-seq, we show that ACF leads to a general increase in Top2 binding across the genome. We map Uls1 binding sites and identify tRNA genes as key regions where Uls1 associates after ACF treatment. Importantly, the presence of Uls1 at these sites prevents ACF-dependent Top2 accumulation. Our data reveal the effect of Top2 poisons on the global Top2 binding landscape and highlights the role of Uls1 in antagonising Top2 function. Remodelling Top2 binding is thus an important new means by which Snf2 enzymes promote genome stability.

## INTRODUCTION

All eukaryotic genomes are organised into chromatin; a complex arrangement of DNA and associated binding proteins. Due to the relative inaccessibility of DNA within chromatin, a universal problem facing eukaryotes is how to access their genetic information. One of the means by which this is achieved is by mechanically altering local chromatin structure through the action of ATP-dependent chromatin remodelling (Snf2) enzymes (1). These proteins are ubiquitous amongst eukaryotes (2) and their influence on chromatin structure means that Snf2 proteins affect all DNA-based transactions such as DNA transcription, replication and repair (3). Underscoring their importance, mutations within human Snf2 proteins cause a range of developmental disorders (4, 5) and SWI/SNF is the most commonly mutated chromatin-regulatory complex in human cancers (6). The majority of Snf2 proteins act by remodelling nucleosomes (1). However, some Snf2 proteins have been shown to act on non-nucleosomal DNA binding proteins such as TBP (7, 8) and Rad51 (9-11). Indeed, for others, their functions remain largely unknown. Here, we use budding yeast to study one such Snf2 factor, *ULS1* and find that its deletion results in hypersensitivity to the Topoisomerase II (Top2) poison acriflavine (ACF).

Top2 is an essential mediator of genome stability due to its ability to disentangle DNA molecules and resolve DNA torsional stress (12). Loss of Top2 causes irreparable defects in cell division whereas blocking Top2 catalytic activity induces massive DNA damage and checkpoint arrest (13). As part of its reaction cycle, Top2 forms a transient protein-DNA adduct termed the cleavage complex (12). If this intermediate is not resolved, it results in the formation of a DNA single-strand or double-strand break next to a covalent Top2-DNA adduct (14, 15); both highly cytotoxic lesions. This enzymatic weakness is targeted by Top2-poisons, which act to stabilise the cleavage complex (15). This is in contrast to the mechanism of Top2 catalytic inhibitors, which do not stabilise cleavage complex formation (16). The ability of Top2 poisons to turn Top2’s enzymatic activity against itself makes them an important class of anti-cancer drugs. However, even in non-cancerous cells, excess topoisomerase activity is potentially dangerous as it increases the probability that some topoisomerase molecules will stall as cleavage complexes. Several endogenous protein inhibitors of topoisomerase activity exist in bacteria (17-19). Therefore, it is perhaps a little surprising that equivalent eukaryotic topoisomerase inhibitors have not previously been described.

We find that Uls1 helps to keep Top2 activity in check by altering its chromatin association. Uls1 binds Top2 via a Top2-interaction domain (amino acids 350-655) and has DNA-stimulated ATPase activity. Both Uls1’s Top2 interaction domain and ATPase activity are essential for its function, consistent with the idea that it remodels chromatin-bound Top2. This is in agreement with a recent report showing that the homolog of Uls1 in the distantly related yeast *Schizosaccharomyces pombe,* can displace Top2 from DNA (20). Moreover, we extend these observations by mapping how Uls1 influences the genome-wide binding distribution of Top2 *in vivo.* Using ChIP-seq, we show that ACF causes a general increase in Top2 binding across the genome, except at Uls1 binding sites. Thus, the presence of Uls1 is sufficient to displace Top2 from chromatin after exposure to ACF. Uls1 binding sites are distributed throughout the genome but, in the presence of ACF, become enriched at tRNA genes. Interestingly, many tRNA genes show a *ULS1*-dependent decrease in Top2 binding after ACF treatment. This reveals unexpected complexity in the function of Uls1 and suggests that targeting related human Snf2 proteins may reduce the toxicity associated with Top2 poisons by sensitising cancers to these drugs (21, 22).

## RESULTS

### Excess Top2 activity is toxic to *uls1Δ* cells

Deletion of *ULS1* does not result in a dramatic growth defect or in sensitivity to a variety of DNA damaging drugs (S1A Fig). This apparent absence of phenotype initially hindered our attempts to understand its function. However, a previous large-scale chemogenetic screen identified ACF as a drug that specifically kills *uls1*Δ yeast (26) and we confirmed the potent toxicity of ACF (Fig 1A). ACF has been described as having antibacterial (27), antimalarial (28) and anti-cancer properties (29). This broad range of activity is likely due to the fact that ACF inhibits type II topoisomerase activity *in vitro* (28, 30). We show that in budding yeast, ACF acts as a Top2 poison rather than as a Top2 catalytic inhibitor. ACF stabilises Top2 cleavage complex formation *in vitro* and ACF toxicity is enhanced by Top2 over-expression *in vivo* (S1B-C Fig) – both hallmarks of Top2 poisons. Our data are consistent with a previous study showing that acriflavine stabilises the formation of type II topoisomerase cleavage complexes within trypanosome mitochondria *in vivo* (31). To explore the pathways targeted by ACF in yeast, we isolated spontaneous ACF suppressor mutants of *uls1*Δ strains in a forward genetic screen. Of the eight independent suppressor colonies tested, all contained single point mutations within *TOP2,* two of which were identified multiple times (Fig 1B). These data show that Top2 is the most significant factor mediating ACF toxicity in yeast.

**Figure 1.**
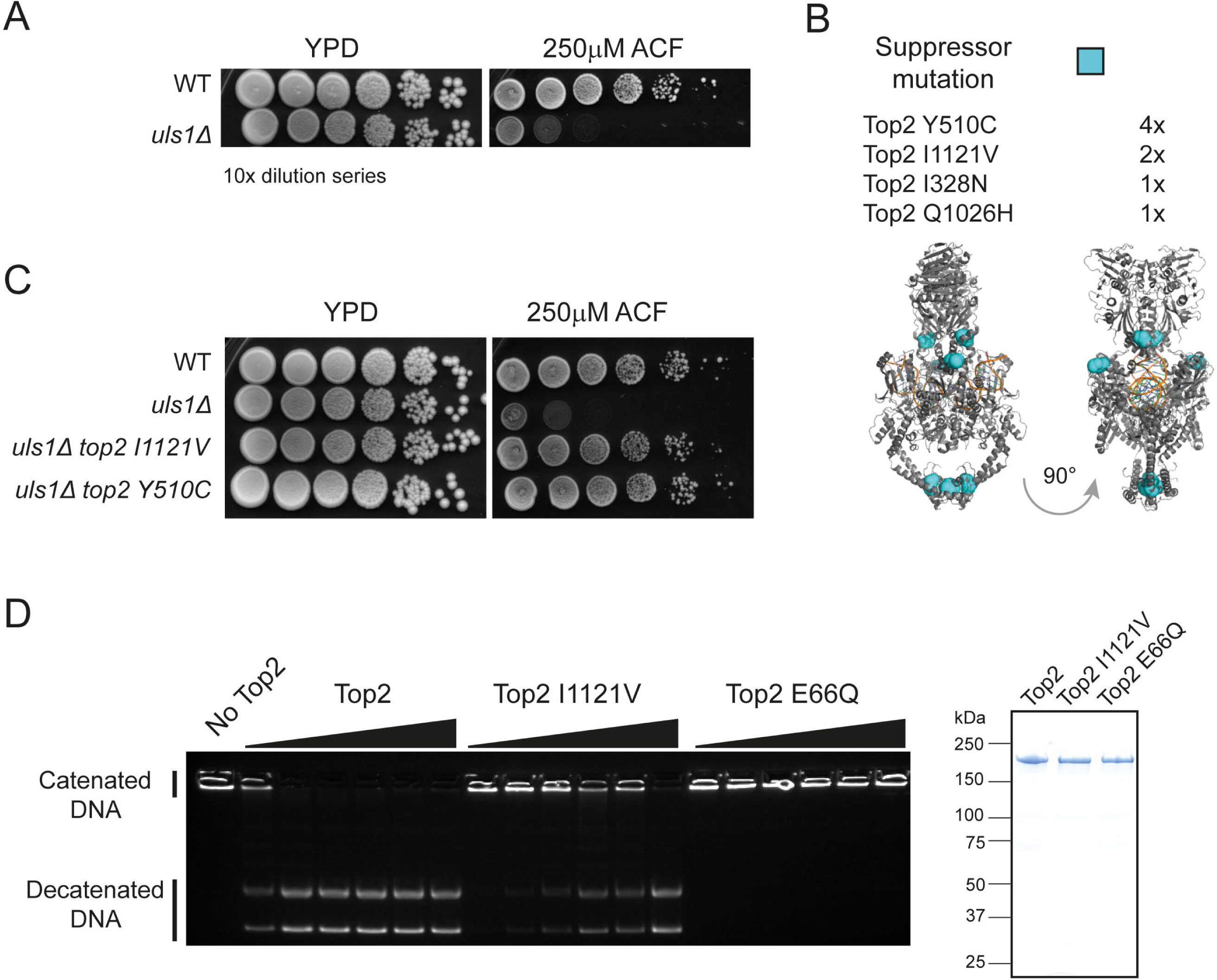
*ULS1* deletion causes sensitivity to ACF due Top2 activity. (A) 10-fold serial dilutions of WT (HFY9) or *uls1*Δ (HFY71) yeast on rich media (YPD) or drug containing plates (ACF). (B) Identification of isolated suppressor mutants and their location within the structure of the Top2 dimer (PDB ID: 4GFH). (C) Top2 point mutations were introduced into independent yeast strains to verify they are causing suppression. *top2 I1121V* (HFY264) and *top2 Y510C* (HFY263) alleles fully supress the ACF sensitivity of *uls1*Δ (HFY71) such that the grow identically to WT (HFY9) on ACF. (D) *in vitro* decatenation assay. 200nM of kinetoplastid DNA was incubated for 30 mins at 30°C with 0, 3, 6, 12, 25, 50 or 100nM Top2 before being run out on a 1% agarose gel. Top2 containing the suppressor mutation I1121V (HFP273) is approximately 16-fold less active than wildtype Top2 (HFP 185) but still has significantly more activity than the ATPase dead Top2 E66Q (HFP271). A Coomassie-stained protein gel on the right illustrates the purity of expressed Top2 constructs.

To test whether *uls1*Δ cells are generally sensitive to Top2 poisons, we additionally tested the Top2 poisons, ellipticine. We find that *ULS1* deletion results in sensitivity to ellipticine but only in a sensitising *rad51*Δ background (S2A Fig). This may reflect subtle differences in their mode of action (32, 33) or in drug uptake. Indeed, Top2 poisons such as etoposide are poorly taken up by yeasts, meaning that drug sensitivity in wildtype cells is typically only observed in genetic backgrounds that contain plasma membrane pump mutations (20, 34). In contrast, we find that ACF uptake from agar plates is very efficient, even in strains without membrane pump mutations. We have taken advantage of this to carry out a genome-wide deletion library screen for ACF sensitivity in an otherwise wildtype yeast background, which will be published elsewhere. We introduced the *TOP2* alleles identified in our ACF suppressor strains into independent yeast strains. This confirmed that the suppression phenotype observed was solely due to mutations in *TOP2* and not of any other factor (Fig 1C). The suppression of the initial *uls1*Δ ACF sensitivity was complete as *uls1*Δ *top2 I1121V* or *uls1*Δ *top2 Y510C* double mutant cells grew indistinguishably from wildtype (Fig 1C). This further reinforces the notion that Top2 is the key target of ACF *in vivo*. Whilst we cannot exclude that ACF affects other cellular pathways, if it does, they do not significantly affect cellular growth or viability.

The ACF suppressor mutations identified did not cluster within the three-dimensional Top2 protein structure (Fig 1B), making it unlikely that they were affecting a protein-protein interaction. Instead, we hypothesized that the suppressor mutations were influencing Top2 catalytic activity. To test this, we purified wildtype and mutant yeast Top2 and carried out *in vitro* decatenation reactions. As seen in Fig 1D, Top2 I1121V was able to unlink the interlocked rings of kinetoplastid DNA, in contrast to the ATPase dead Top2 E66Q allele. However, Top2 I1121V was approximately 16-fold less active than wildtype. These data are consistent with ACF acting as a Top2 poison as reduced Top2 enzymatic activity results in lower drug toxicity. Consequently, the most likely reason that *uls1*Δ cells are more sensitive to ACF than wildtype is that they have increased Top2 activity. This antagonism between Uls1 and Top2 is not just drug dependent as overexpression of Top2 is toxic to *uls1*Δ yeast, even in the absence of ACF (S1C Fig).

### Amino acids 350-650 within Uls1 mediate physical interaction with Top2

Having established a genetic interaction between Top2 and Uls1, we asked the question whether these two proteins interact physically. Using a yeast 2-hybrid (Y2H) assay, we detected weak but reproducible binding between full-length Uls1 and full-length Top2 *in vivo*. Furthermore, we could narrow down the region of Uls1 required for Top2 interaction to fragment 350-655 (Fig 2A). To verify that the Uls1-Top2 binding interaction observed was direct, we assayed their ability to interact *in vitro.* Using purified proteins, we confirmed that Uls1 fragment 350-655 binds to Top2 *in vitro* (Fig 2B). This region of Uls1 contains several putative SUMO-interaction motifs (SIMs) (35) and is able to bind SUMO by Y2H assay (S3A Fig). Moreover, Top2 can be sumoylated *in vivo* (36). However, the purified Top2 used in our *in vitro* binding assays had no detectable sumoylation, as determined by mass spectrometry (data not shown). Therefore, Uls1 binding to Top2 is unlikely to require Top2 sumoylation, although it might be enhanced by it.

**Figure 2.**
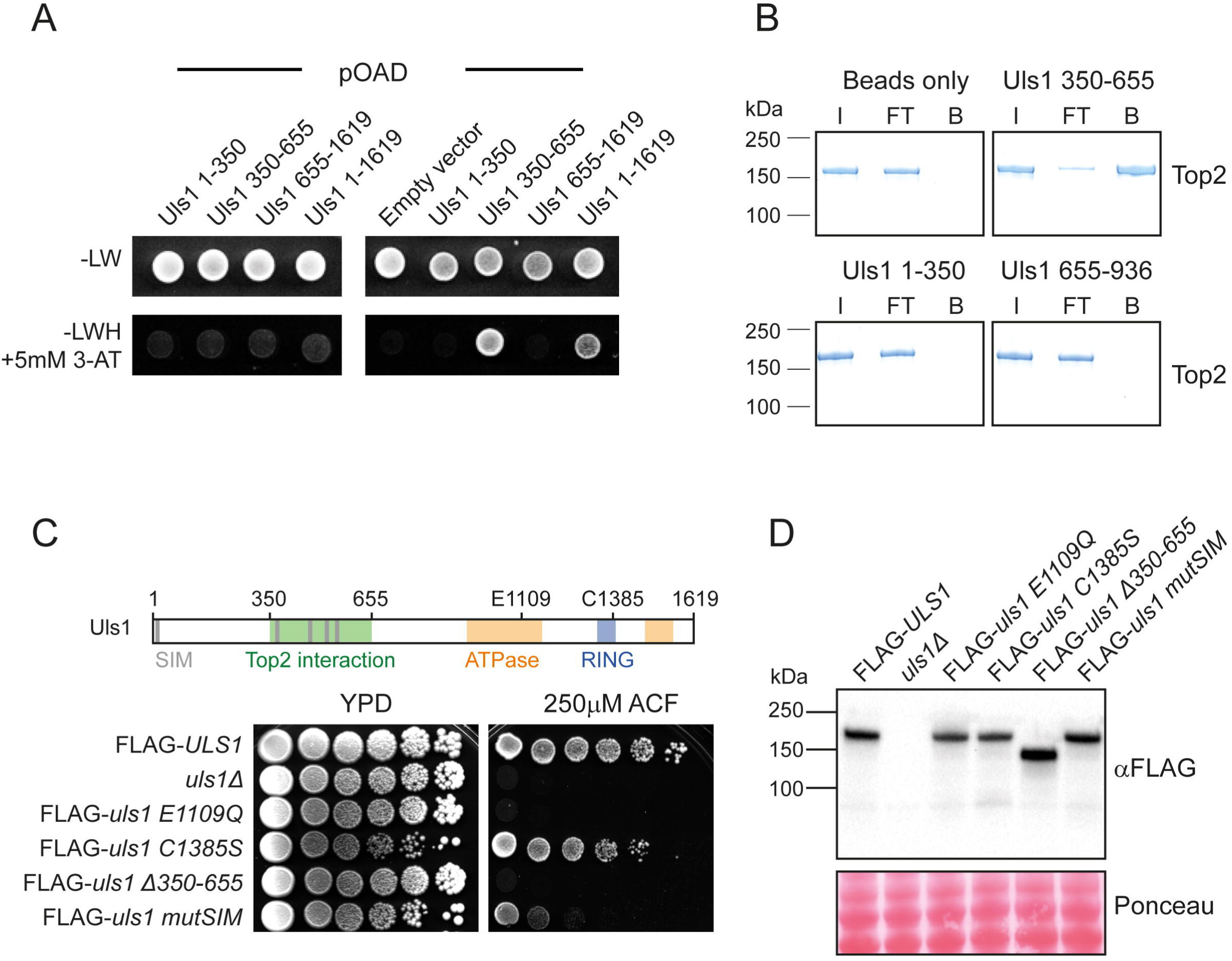
Physical interaction of Uls1 and Top2 is important for Uls1 function. (A) Yeast 2-hybrid assay. Yeast containing the indicated combination of Gal4 activator domain (pOAD) and Gal4 binding domain (pOBD) plasmids were grown on control (-LW) plates and assay (-LWH with 5mM 3-Amino-1,2,4-triazole) plates. Full length Uls1 (HFP136) and Uls1 350-655 (HFP133) interact with Top2 (HFP 185) but not the empty vector control (HFP122). In contrast, Uls1 fragments 1-350 (HFP193) and 655-1619 (HFP134) do not bind Top2. (B) *in vitro* pulldown of full length Top2 with the indicated fragments of Uls1 bound to agarose beads showing input (I), flow-through (FT) and bound (B) fractions. (C) Diagram of Uls1 domain architecture. Serial dilutions of the indicated genotypes were assayed for viability on 250μM ACF. Mutation of *ULS1* ATPase function (*uls1 E1109Q* - HFY275) or deletion of its Top2 interaction domain (*uls1* Δ*350-655* - HFY225) mimics *uls1*Δ (HFY71). In contrast, mutation of ULS1’s RING finger (*uls1 C1385S* - HFY230) has hardly any effect on ACF sensitivity whereas mutation of its five putative SIMs (HFY261) has a moderate effect on ACF sensitivity. (D) Western blot of the same constructs used in (C) indicating equivalent expression levels. Ponceau-stained membrane is used a loading control.

To assess the functional significance of Uls1-Top2 interaction, we introduced a range of mutations into the endogenous *ULS1* gene and FLAG-tagged it to monitor its expression level. Strikingly, deletion of the Top2 interaction domain, *uls1* Δ*350-655*, mimicked complete loss of *ULS1* (Fig 2C). In contrast, mutating all predicted SIMs in Uls1 resulted in only moderate ACF sensitivity. These data show that Top2 interaction is essential for Uls1 activity whereas SUMO-binding merely promotes it. As expected for a Snf2-family enzyme, mutating the Walker B motif (E1109Q) within the ATPase domain of Uls1 completely inactivated its function. However, mutating Uls1’s RING domain (C1385S) had no significant effect (Fig 2C). It is important to note that none of the phenotypes observed are due to altered Uls1 protein levels (Fig 2D). Uls1 has previously been proposed to act as a SUMO-targeted Ubiquitin Ligase (STUbL), with SUMO-targeting being mediated via its SIMs and the RING domain acting as an E3 Ubiquitin ligase (35). However, in the context of ACF resistance, we see that Uls1’s RING domain is dispensable, and that SIMs play an important but non-essential role. Therefore, it appears unlikely that Uls1 is acting as a STUbL on Top2 and indeed, Top2 protein levels do not change significantly in *uls1*Δ strains (S3B Fig).

### Uls1 has weak DNA stimulated ATPase activity

ATP-hydrolysis is an essential feature of all Snf2 proteins (1). To characterise Uls1’s ATPase activity, we attempted to purify the full-length protein from yeast. However, Uls1 is a large (184kDa), low abundance protein and overexpressing it in yeast or *Sf9* insect cells gave very poor yields. We noticed that deleting the first 349 amino acids of Uls1 resulted in a significant increase in yeast expression (data not shown). Amino acids 327-350 contain a predicted nuclear localisation signal (NLS). However, in terms of catalytic function, the Uls1 Δ1-349 protein is fully active (S3C Fig) and therefore suitable for biochemical characterisation.

Uls1 ATP hydrolysis was monitored via a coupled enzymatic reaction utilizing pyruvate kinase and lactate dehydrogenase to oxidise NADH (25) (Fig 3A). We find that Uls1 displayed weak DNA-stimulated ATPase activity (Fig 3B). This ATPase activity is due to Uls1 and not a contaminating protein as it was abolished in an ATPase mutant (E1109Q) version of Uls1 (Fig 3B-C). We also tested whether Uls1’s ATPase activity would be activated by Top2 *in vitro*. However, we were unable to detect any measureable Uls1-dependent increase in ATPase activity in the presence of Top2 (S4 Fig). This was also true if we used a version of Top2 with a 5xSUMO tag on its C-terminus to mimic endogenous sumoylation (data not shown). These assays were hampered by the very low amounts of Uls1 that we were able to purify. It is possible that the concentrations of Uls1 used may be below its association constant for Top2 or that we have not used appropriate reaction conditions, making it difficult to draw strong conclusions from these experiments. However, importantly, we have been able to show that purified Uls1 has DNA-stimulated ATPase activity. To the best of our knowledge, all Snf2-family enzymes tested have shown DNA-stimulated ATPase activity *in vitro* as they all act on DNA-bound substrates *in vivo (8, 37-39)*. Therefore, Uls1 behaves functionally as a *bone fide* Snf2 protein.

**Figure 3.**
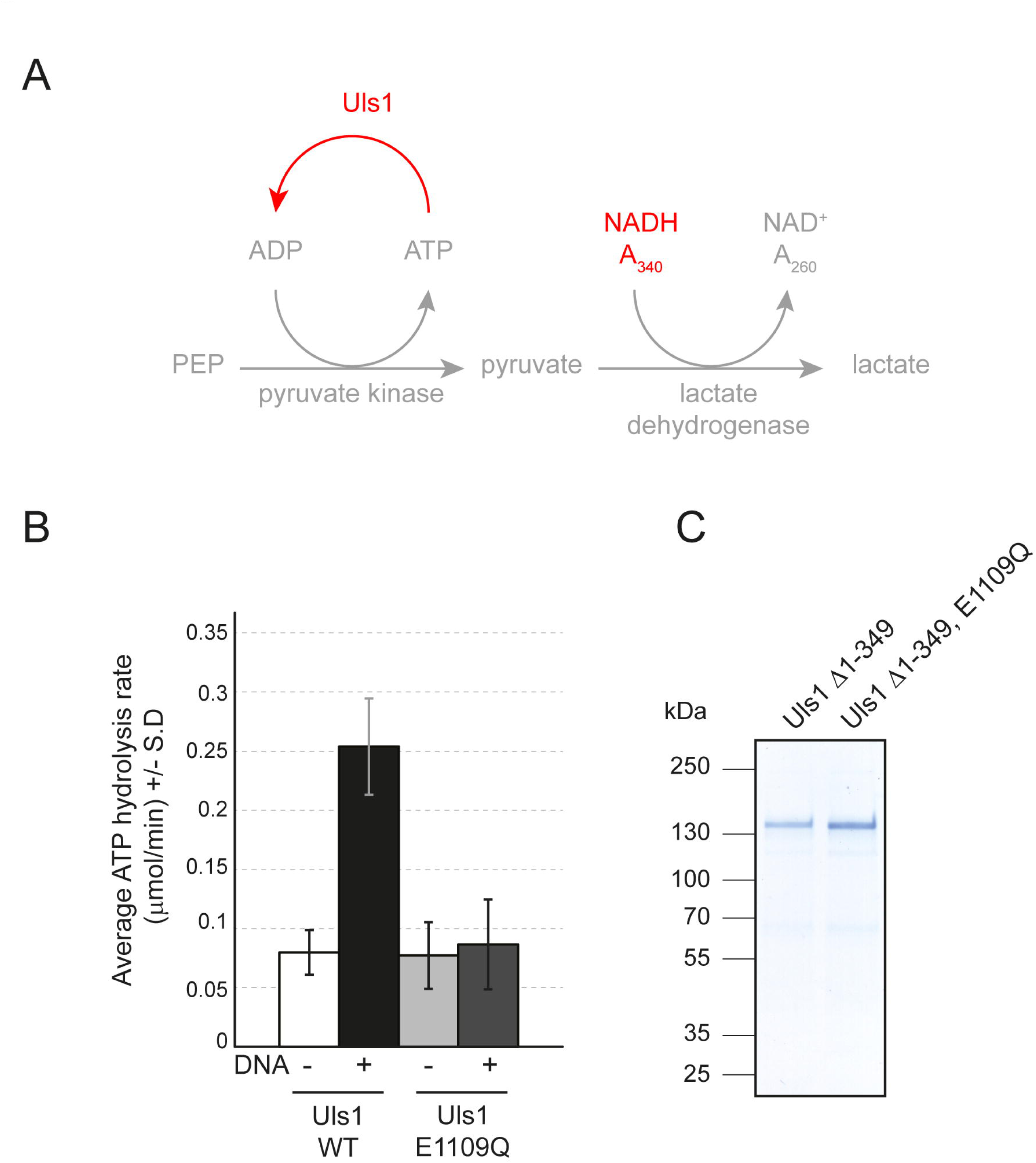
Uls1 has DNA-stimulated ATPase activity. (A) Scheme of the coupled ATPase assay used, reactions were carried out at 30°C and A_340_ measurements taken every 10s for 30 mins. (B) ATP hydrolysis rates for the indicated proteins. The graph shows the average +/- the standard deviation of three independent experiments. 15nM Uls1 was incubated with or without 100μM salmon sperm DNA. (C) A Coomassie-stained protein gel on the right illustrates the purity of the purified Uls1 constructs Uls1 Δ1-349 (HFP385) and Uls1 Δ1-349, E1109Q (HFP404).

### Deletion of *ULS1* results in a global increase in acriflavine-stabilized Top2 on DNA

Because of the antagonistic relationship between Uls1 and Top2 activity (Fig 1C and Fig 1E), we decided to test whether Uls1 influenced Top2 localisation *in vivo*. To this end, we performed ChIP-seq on strains with an extra HA-tagged copy of *TOP2* under the control of its endogenous promoter in wildtype (HFY250) and *uls1*Δ (HFY252) cells both in the presence and absence of 250μM ACF. These strains were used as they show the expected ACF sensitivity in a *uls1*Δ background. In contrast, a *uls1*Δ strain where only the endogenous copy of *TOP2* is HA-tagged has suppressed ACF sensitivity (S3B Fig). Four independent ChIP replicates of each condition were pooled to form two DNA sequencing replicates which were aligned to the W303 genome reference (40) using BWA (41) and subjected to automated peak calling by MACS2 software (42). As expected of a Top2 poison, we saw that ACF caused an increase in the number of Top2 peaks called (S5A Fig). Importantly, ACF also caused a significant increase in the intensity of Top2 peaks. Due to the large number of data points involved, statistical significance was assessed using Cohen’s d (*d*), which measures effect sizes based on the difference between two means. Cohen’s d values of 0.2, 0.5 or 0.8 typically denote a small, medium or large effect respectively (43). By performing a pairwise comparison of common peaks, we saw that the addition of ACF resulted in a modest increase (*d* = 0.49) in the average Top2 peak intensity in wildtype cells (Fig 4A). Strikingly, the increase in Top2 peak intensity after ACF treatment in a *uls1*Δ strain (Fig 4B) was much more pronounced (*d* = 1.56). By directly comparing common Top2 peaks between wildtype and *uls1*Δ cells exposed to ACF, we could confirm that significantly more Top2 (*d* = 0.62) becomes DNA-bound in *uls1*Δ cells compared to wildtype (Fig 4C). These data explain the genetic interactions we had seen and suggest that *uls1*Δ cells exposed to ACF die because an excessive amount of Top2 becomes bound to chromatin. Top2 ChIP qPCR in strains where only the endogenous *TOP2* gene is HA-tagged confirmed the trends we were seeing via ChIP-seq (S5B Fig). These data also suggest that *TOP2* copy number does not bias ACF-dependent changes in Top2 chromatin association.

**Figure 4.**
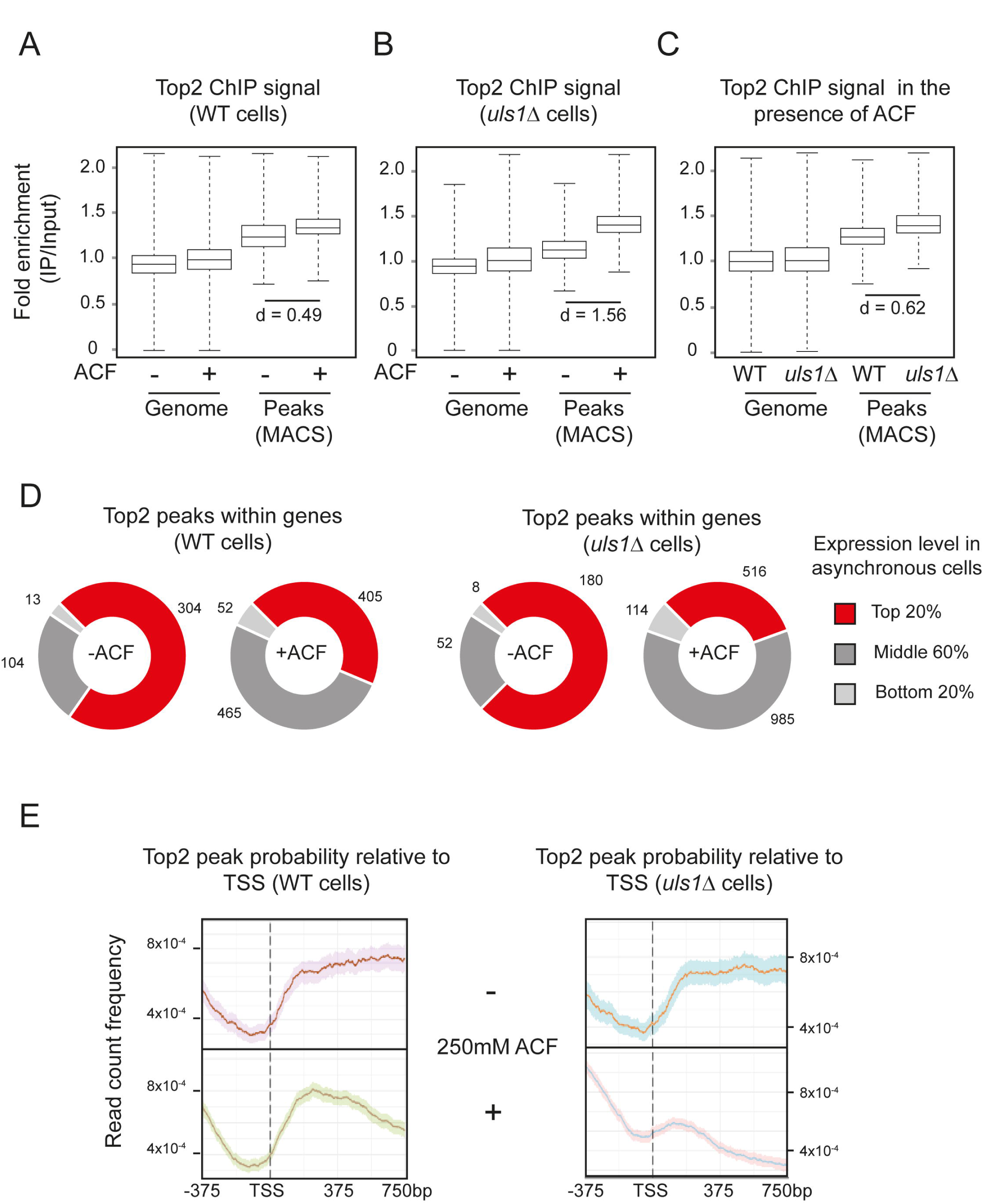
Uls1 controls Top2 chromatin binding in the presence of ACF. (A) Pairwise comparison of the average ChIP enrichment across all mapped reads (Genome) and specifically within common regions called as peaks by MACS2 (Peaks) in wildtype cells (HFY250) both in the presence or absence of 250μM ACF. Top2 peaks become significantly more intense when ACF is added, Cohen’s *d* = 0.49. (B) The same as in A, except in *uls1*Δ cells (HFY252) showing that the effect of ACF is exacerbated, Cohen’s *d* = 1.56. (C) Pairwise comparison of the average ChIP enrichment in the presence of 250μM ACF. Comparing common ACF-dependent peaks between wildtype (HFY250) and *uls1*Δ (HFY252) cells indicates that there is significantly more Top2 bound in *uls1*Δ, Cohen’s *d* = 0.62. (D) Association of Top2 peaks within genes and the expression level of those genes in asynchronous culture under exponential growth. Expression data was taken from (45) and the number of peaks within each group is displayed next to the graph. (E) Normalised Top2 peak probability relative to the TSS of RNAP II transcripts in wildtype (HFY250) or *uls1*Δ (HFY252) cells in the presence or absence of ACF. The solid line displays the average with 95% confidence intervals indicated by the shaded area.

Top2 is known to be associated with ongoing transcription (44). Consistent with this, we find that when Top2 peaks are near genes, these are highly expressed under conditions of exponential growth (45) (Fig 4D). The addition of ACF results in an overall increase in Top2 peak number as well as the distribution of peaks becoming much less biased towards highly expressed genes. This shows that ACF-dependent Top2 peaks are associated with genes but are largely uncoupled from their initial transcription level in unperturbed cells. Interestingly, a similar trend in seen with human cells, where TOP2A-dependent cleavage complex formation within protein coding genes is independent of transcription level (46). By plotting Top2 peak probability relative to the transcription start site (TSS) of the ‘average’ RNA Pol II transcribed gene, we find that Top2 is more likely bound within gene bodies both in WT and *uls1*Δ cells (Fig 4E). Interestingly, this pattern is largely unchanged when WT cells are exposed to ACF. In contrast, *uls1*Δ cells exposed to ACF display a dramatic change such that Top2 peaks are now more likely to be found upstream of the TSS within intergenic regions rather than within coding sequences (Fig 4E). Therefore, *uls1*Δ cells exposed to ACF not only have increased levels of Top2 bound to DNA but its distribution across genes becomes markedly disrupted.

### Uls1-bound regions do not accumulate Top2 after exposure to ACF

We decided to map Uls1 binding sites by performing ChIP-seq on a FLAG-tagged Uls1 strain in the presence and absence of ACF. We used 100μM ACF as Uls1 activity is essential at this concentration (S2B-C Fig) and higher drug concentrations disrupted Uls1 pulldown (data not shown). Overall, there was a slight decrease in the number of unique Uls1 peaks in the presence of ACF and no significant change (*d* = 0.05) in the average Uls1 peak intensity (Fig 5A). This indicates that the absolute level of chromatin-bound Uls1 remains largely unchanged by ACF. However, ACF does re-distribute Uls1 to regions upstream of RNA Pol II genes (Fig 5B).

**Figure 5.**
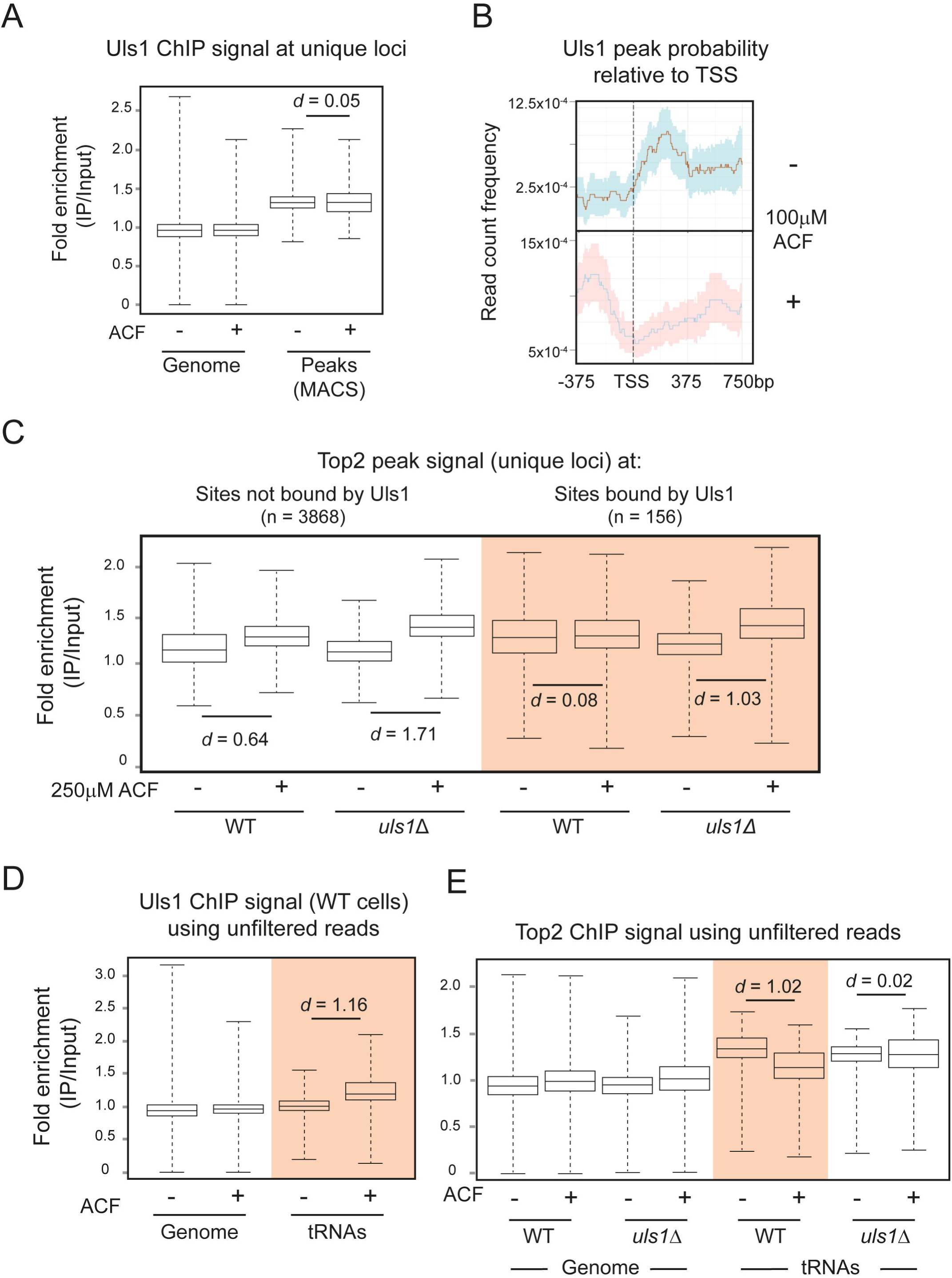
Uls1 binding sites do not accumulate Top2 in the presence of ACF. (A) Pairwise comparison of the average Uls1 ChIP enrichment (HFY176) across all mapped reads (Genome) and specifically within peak regions +/- 100μM ACF. The level of Uls1 chromatin binding is independent of ACF. (B) Normalised Uls1 peak probability relative to the TSS of RNA Pol II transcribed genes in the presence or absence of ACF. The solid line displays the average with 95% confidence intervals indicated by the shaded area. (C) Comparison of the average Top2 ChIP enrichment (using filtered reads) between regions that are either bound or unbound by Uls1 +/- 250μM ACF. In contrast to unbound sites, Uls1 binding sites do not accumulate Top2 in the presence of ACF. This effect is *ULS1* dependent. (D) Pairwise comparison of the average Uls1 ChIP enrichment using unfiltered reads across the genome and specifically within tRNA genes +/- 100μM ACF. Uls1 becomes enriched at tRNA genes in the presence of ACF, Cohen’s *d* = 1.16. (E) Same as (D) except looking at Top2 ChIP. ACF causes loss of Top2 from tRNA genes, which is *ULS1* dependent.

To test our hypothesis that Uls1 was directly influencing Top2 *in vivo*, we compared the behavior of Top2 peaks that either did or did not overlap with Uls1 peaks. At Top2 peaks that do not overlap with Uls1, ACF caused an increase in the amount of Top2 bound to DNA and this effect was exacerbated in *uls1*Δ cells (Fig 5C). This was similar to the trends we had observed previously (Fig 4A-B). However, strikingly, at Top2 peaks that overlap with Uls1, ACF did not cause any significant increase (*d* = 0.08) in Top2 levels. Importantly, in *uls1*Δ cells, the addition of ACF resulted in an increase (*d* = 1.03) in Top2 binding at these sites (Fig 5C). These data support the model that Uls1 acts to remove Top2 trapped on chromatin by ACF.

When we looked specifically for ACF-dependent Uls1 binding sites, tRNA genes stood out. These accounted for 21% of all Uls1 peaks in the presence of ACF, but only 4% in untreated cells (S6A Fig). Most tRNA genes are duplicated in the yeast genome, with some present in as many as 16 copies per cell (47). Our standard bioinformatic analysis filters out sequence reads that map to multiple genomic locations. Therefore, due to their repetitive nature, we might be missing relevant information. By analysing unfiltered sequence reads, we see that Uls1 signal at tRNAs increases significantly (*d* = 1.16) after the addition of ACF (Fig 5D). Indeed, after looking at other repetitive loci (telomeres, rDNA and Ty retrotransposons), tRNA genes are the only regions where Uls1 signal increases significantly after ACF treatment (S6B Fig). Importantly, we also observe an antagonistic relationship between Uls1 and Top2 at tRNA genes. ACF caused a significant decrease (*d* = 1.02) in Top2 signal at tRNA genes which was *ULS1*-dependent (Fig 5E). Thus, the presence of Uls1 prevents ACF-dependent Top2 accumulation at tRNA genes as it does at other genomic loci.

## DISCUSSION

We show here that Uls1 can suppress Top2 activity by removing Top2 that becomes chromatin-bound when cells are exposed to the Top2 poison ACF. Our ChIP procedure cannot differentiate between a true Top2 cleavage complex and Top2 that is non-covalently bound to DNA. However, the distribution of ACF-dependent Top2 peaks in yeast are consistent with the behaviour of *bona fide* TOP2A cleavage complexes in human cells (46) as both are independent of transcription level. This suggests that Top2 poisons are opportunistic in their mode of action and will trap Top2 molecules wherever they are found.

Although ACF leads to a general increase in Top2 binding to chromatin, there are a few regions including ribosomal protein genes (S5C Fig), tRNA genes and the rDNA locus (S6C Fig) where ACF resulted in a decrease in the amount of Top2 bound. It is not immediately clear why ACF should cause less Top2 to be DNA-bound at these sites. However, it is possible that stalled Top2 at these highly transcribed genes is more easily detected and targeted for degradation. Indeed, one of the main mechanisms of recognising Top2 adducts is via collision with the transcription machinery (48). Overall, the effects of ACF become exacerbated when *ULS1* is deleted: more Top2 peaks are found and their signal intensity is higher, consistent with more Top2 becoming chromatin-bound. We see that Uls1 tends to bind close the to 5’ end of RNA Pol II gene coding regions, in agreement with what has been observed for several other Snf2 proteins (49, 50). In the presence of ACF, a significant fraction of Uls1 relocalises to tRNA genes. Importantly, at Uls1 peaks, there is no ACF-dependent increase in chromatin-bound Top2, suggesting that Uls1 removes Top2 from DNA (Fig 6A-B).

**Figure 6.**
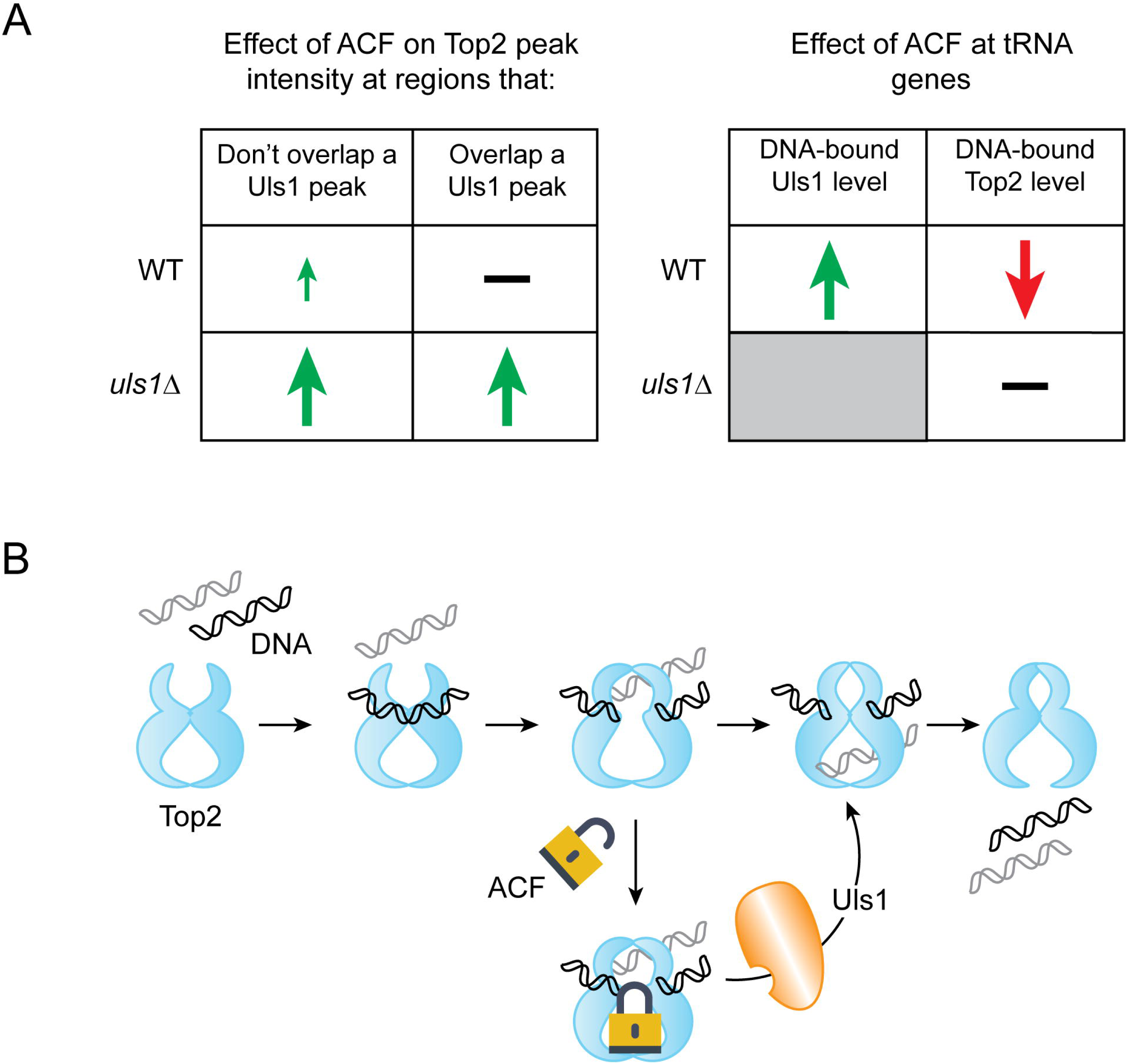
Model of how Uls1 and acriflavine influence Top2 DNA binding. (A) Summary of ChIP data describing how Uls1 antagonises the ACF-dependent increase in Top2 binding throughout the genome. (B) Model of how Uls1 might remodel a Top2 cleavage complex by promoting DNA-stimulated Top2 ATPase activity leading to movement of the transfer DNA (grey) and resolution of the Top2-DNA bonds within the guide DNA (black).

We do not always see a direct anti-correlation between DNA-bound Top2 and Uls1. This may, in part, be because there is almost 30 times more Top2 than Uls1 in a yeast cell (51). Consequently, deletion of *ULS1* results in ACF-dependent changes in Top2 binding at far more sites than we see Uls1 binding to. We cannot exclude that some of these effects are indirect. Moreover, Uls1-Top2 interaction may be dynamic and so Uls1 may only interact transiently at any given site before dissociating away to bind another region. This is not atypical for Snf2 proteins whose ATPase activity can influence substrate binding (52, 53).

The precise mechanism by which Uls1 remodels Top2 to release it from the cleavage complex is uncertain. We see that Uls1 function is completely dependent on its ATPase activity, partially dependent on SUMO interaction and independent of its RING domain. This suggests that, at least within this context, Uls1 is not acting as a STUbL to degrade proteins (35). Snf2 proteins are known to translocate along DNA in an ATP-dependent manner (54). We therefore speculate that Uls1 may use its DNA translocase activity to alter Top2-DNA interactions. This may displace Top2 from DNA or potentially alter the precise orientation of DNA within a Top2 cleavage complex and so stimulate Top2’s intrinsic ATPase activity to release itself from DNA (23). It is not clear at this stage why Uls1 is recruited to tRNA genes to remodel Top2. There is very little published literature linking tRNA genes with Top2. However, topoisomerase activity appears to be largely dispensable for tRNA transcription in yeast (55). Therefore, it is possible that Uls1 is being recruited to tRNA genes to deal with stalled Top2 not because of an effect on tRNA expression but because of replication fork arrest, which occurs primarily at tRNA genes in yeast (56).

Utilising Uls1 to remodel trapped Top2 may be particularly important in lower eukaryotes as they lack the pathway used by mammals to cleave the 5’-phosphotyrosyl bond within covalent Top2-DNA complexes (57, 58). It remains to be seen whether mammalian homologs of Uls1 can carry out analogous Top2 remodelling reactions. If so, it opens up the possibility of targeting these Snf2 proteins in combination with Top2 poison treatment to potentiate anticancer therapies.

## MATERIAL AND METHODS

### Yeast strains

A full strain list (S1 Text) and plasmid list (S2 Text) can be found in supplementary information.

### Protein expression and purification

Full length Top2 (HFP185 - a gift from J. Berger) and mutants E66Q or I1121V (HFP271, HFP273) were expressed as previously described (23). For WT and E1109Q Uls1 expression (HFP 385, HFP404), plasmids were transformed into HFY155. 6L of YPLG media was inoculated (1:10 ratio) with a saturated overnight culture (SC-URA) and incubated at 30°C for 16 hours. Protein expression was induced by the addition of 2% galactose (final) and the culture harvested after 6-hour cultivation at 30°C. A cryogenic grinder was used to disintegrate yeast cells. The powder was diluted in Lysis buffer (50mM HEPES; pH 7.4, 500mM NaCl, 10mM imidazole, 10% glycerol, 0.5% Triton X-100 and EDTA-free protease inhibitors (Roche)) and spun at 35,000g for 1 hour at 4°C. The supernatant was incubated for 30 mins with TALON resin (Clontech), washed extensively with TALON wash buffer (50mM HEPES; pH 7.4, 500mM NaCl, 10mM imidazole, 10% glycerol) and eluted with TALON elution buffer (50mM HEPES; pH 7.4, 500mM NaCl, 200mM imidazole, 10% glycerol). The eluted protein was loaded onto a Strep-Tactin XT column 1 ml (IBA), washed with Strep-Tactin wash buffer (50mM HEPES; pH 8.0, 200mM NaCl, 10% glycerol) and eluted by Strep-Tactin elution buffer (50mM HEPES; pH 8.0, 200mM NaCl, 10% glycerol, 50mM Biotin). The eluted protein was concentrated using a 10 kDa MWCO Amicon spin column, frozen in liquid N_2_ and stored in small aliquots at -80°C.

### *[H2]in vitro* protein interaction assay

Top2 (prey) was expressed and purified as described above. To obtain the bait protein, BL21(DE3)RIL *E. coli* was transformed with the relevant plasmids (HFP219, HFP221, HFP222). The cells were grown in TB medium at 37°C until OD_600_ = 0.4-0.6. Expression was induced with 0.5mM IPTG and left for 16-18 hours at 16°C. The pellets were resuspended in Lysis buffer, sonicated and centrifuged at 4°C, 20,000g for 1 hour. The supernatants were added onto TALON resin (Clontech) and incubated at 4°C for 40 min. The resins were washed with TALON wash buffer and eluted with TALON elution buffer. Approximately 0.1 mg of bait protein was pre-bound with 80μl of Strep-Tactin superflow (IBA) beads and washed with Pulldown buffer (25mM HEPES; pH 7.5, 150mM KCl, 3mM MgCl2, 5% glycerol, 1mM DTT, 0.1% NP-40). 200μl of the prey protein (0.1 mg/ml) was added to the beads and incubated together with the bait or empty beads for 1 hour at 4°C. Then the beads were washed three times with Pulldown buffer and 20μl of 5 × SDS-Sample buffer was added directly to the beads and boiled together with input and flowthrough fractions. The bound fraction is approximately 20 × more concentrated than input and flow through fractions.

### Topoisomerase activity assays

Decatenation assays were performed using a Topoisomerase II Assay kit (TopoGEN, TG1001-1) except with yeast Top2. The reaction was incubated for 30 minutes at 30 °C and terminated by the addition of 5x Stop buffer. Samples were loaded onto a 1% agarose gel containing 0.5 μg/ml of ethidium bromide and run for 1 hour at 4 V/cm. Plasmid linearization assays was performed as described previously (24) with minor modifications. The reaction volume was 20μl. 2μl of 1μM Top2 (homodimer) was added into the tube containing 5 nM pUC19 vector (166.7ng), +/- etoposide or acriflavine in appropriate concentration and 2μl of 10× reaction buffer (500mM Tris·Cl; pH 8, 100mM MgCl2, 5mM dithiothreitol, 1.5M NaCl, 300μg/ml BSA). The mixed reaction was incubated at 30°C for 15 min.

The reaction was terminated by adding 2μl of 10% SDS. Then 1.5μl of 250mM EDTA and 2μl of 1mg/ml proteinase K was added, incubating for 2 hours at 50°C. Samples were loaded on a 1 % agarose gel containing 0.5μg/ml EtBr with electrophoresis carried out for 3hr at 4 V/cm.

### ATPase assay

An enzyme-coupled ATPase assay based on hydrolysis of ATP coupled to oxidation of NADH was used to measure the protein ATPase activity (25). 15nM Uls1 and/or 50μM homodimeric Top2 alone or with 100μM DNA (purified sheared salmon-sperm DNA, Invitrogen) were mixed together in a buffer containing 50mM Tris.HCl; pH 7.9, 100mM KCl, 8mM MgCl2, 5mM beta-mercaptoethanol, 200ug/ml BSA, 2mM Phospho(enol)pyruvate, 280μM NADH (Sigma, N7410), 0.5mM ATP and 1ul of pyruvate kinase/lactate dehydrogenase mix (Sigma, P0294). The reactions were performed in 100μl reaction volume in a 96 well-plate at 30 °C. The oxidation of NADH to NAD+ was monitored by measuring of the fluorescence (excitation - 340 nm, Emission - 440m) every 30s for 30 min using a Spectramax Gemini XPS microplate reader. Titration of increasing concentration on NADH was used to obtain a standard curve for each measurement. The background signal was subtracted from each sample before plotting the results into the graph.

### Chromatin Immunoprecipitation

Cells were grown to OD_600_ 0.6, split in two and then incubated with or without ACF for two hours. Yeast in ACF containing media were spun and re-suspended in an equivalent volume of fresh YPD before crosslinking with 1% formaldehyde for 10 minutes and quenching with 140mM glycine.

Yeast were disrupted using homogenization beads (0.5mm diameter, Thistle Scientific 11079105) in 200μl lysis buffer (50mM HEPES pH 7.5, 140mM NaCl, 1mM EDTA, 1% Triton X-100, 0.1% sodium deoxycholate, protease inhibitors). They were bead beaten in a FastPrep disruptor for 5 × 30 seconds at power setting 6.5, with cooling on ice between each cycle. Lysates were diluted in a further 300μl lysis buffer and spun for 15 minutes at 15,000 rpm at 4°C. The pellet was resuspended in 300μl lysis buffer in a 1.5ml Bioruptor tube (Diagenode, C30010016) and chromatin sheared using a Bioruptor Pico, 10 cycles of 30s on/off (DNA should be sheared to fragments of 250-500bp). This was centrifuged at 8,000 rpm for 5 minutes at 4°C and the supernatant used for ChIP.

25μl magnetic Protein A/G beads (Fisher, 11844554) and 1μg antibody (anti-FLAG: Sigma, F3165 or anti-HA: Roche, clone 3F10, ROAHAHA) per test condition are added to 500μl 5mg/ml PBS-BSA which is rotated for 1 hour at 4°C. This was washed with lysis buffer and then incubated with ChIP extract for 3 hours at 4°C. Beads are washed twice with lysis buffer for 5 minutes and then twice with wash buffer (100mM Tris pH 8, 250mM NaCl, 0.5% NP-40, 0.5% sodium deoxycholate, 1mM EDTA, protease inhibitors) before elution in 60μl TE, 1% SDS at 65°C for 15 minutes.

To prepare protein samples for gel-electrophoresis, samples are un-crosslinked by boiling at 95°C for 15 minutes before loading onto the gel. To prepare DNA for purification, 1% SDS is added to input, 0.5μl RNase A (10mg/ml) is added to both input and IP DNA, and both samples are un-crosslinked overnight at 65°C in a PCR machine. 0.5μl Proteinase K (20mg/ml) is added after uncrosslinking and samples incubated for 1 hour at 65°C. DNA was purified using Qaigen QIAquick PCR purification kit (Qiagen, 28106) as per specifications, eluting in 50μl H2O.

### DNA sequencing and ChIP-seq analysis

A detailed description of library preparation and bioinformatics analysis (S3 Text) can be found in supplementary information.

## ACKNOWLEDGEMENTS

We’d like to thank Susan M. Gasser (FMI, Basel), in whose lab this project originated, the St Andrews Bioinformatics Unit, Andy Cassidy (Tayside Centre for Genomic Analysis) and David Dickerson (Dundee) for support with the generation and analysis of ChIP-seq data. Michael Lisby (Copenhagen) carried out a genetic screen which informed this project and will be published elsewhere. Michaela Dermendjieva made tagged strains as part of a BBSRC/EASTBIO Research Experience Placement. We would also like to thank Stuart MacNeill, Malcolm White, Kazunori Tomita and Vincent Dion for critical comments and suggestions.

---

**Supplementary figure S1. Acriflavine is a Top2 poison.**

(A) Deletion of *ULS1* does not cause sensitivity to the replication inhibitor, Hydroxyurea (HU), the DNA SSB and DSB forming drug Zeocin or the Top1 poison Camptothecin. (B) *in vitro* cleavage assay. 200nM supercoiled pUC18 plasmid DNA (scDNA) was incubated for 30mins at 30°C with the indicated amounts of Top2 and either etoposide or ACF. Addition of ACF induced DNA cleavage, seen by the appearance of linear DNA, at lower concentrations than the positive control Top2 poison, etoposide. (C) 10-fold serial dilution of yeast containing either an empty vector or a vector driving expression of Top2 (HFY185) under control of the *GAL1* promoter. Overexpression of Top2 is toxic to *uls1*Δ cells and is synergistically lethal with ACF.

**Supplementary figure S2. Deletion of *ULS1* sensitises yeast to ellipticine and ACF activates the DNA damage checkpoint**

(A) 10-fold serial dilution of yeast showing that *ULS1* deletion causes sensitivity to the Top2 poison Ellipticine but only in a sensitising background. A *uls1*Δ, *rad51*Δ double mutant strain (HFY33) is significantly more sensitive than a single *rad51*Δ strain (HFY27). (B) 100μM ACF is toxic to *uls1*Δ cells. (C) 100μM ACF is sufficient to induce robust activation of the DNA damage checkpoint in *uls1*Δ yeast as visualised by Rad53 phospo-shift using an anti-Rad53 antibody (Abcam 104232).

**Supplementary figure S3. Deletion of *ULS1* does not alter Top2 protein levels**

(A) Yeast 2-hybrid assay showing that full-length Uls1 (HFP136) and Uls1 35-655 (HFP133) can interact with Smt3 (yeast SUMO) *in vivo* (HFP288). (B) Top panel shows Top2 protein levels as measured by Western blot using anti-Top2 (TopoGEN TG2014), or anti-HA (Roche ROAHAHA) antibodies with an anti-Tubulin (Sigma T5168) loading control. Top2 protein levels are comparable between congenic wildtype and *uls1*Δ yeast (HFY9 with HFY71, HFY294 and HFY295 with HFY297 and HFY250 with HFY252). The bottom panel illustrates that HA tagging the endogenous *TOP2* locus (HFY297) suppresses ACF sensitivity in contrast to introducing an extra HA-tagged copy of *TOP2* (HFY252). (C) 10-fold serial dilutions of the indicated genotypes showing that Uls1 needs to be nuclear for its function and that the first 349 amino acids contain a nuclear localisation sequence (NLS). *uls1*Δ *1-349* (HFY234) phenocopies *uls1*Δ (HFY71). However, its function is fully rescued by addition of an SV40 NLS (HFY281).

**Supplementary figure S4. Top2 does not stimulate Uls1’s ATPase activity**

(A) ATP hydrolysis rates for the indicated proteins. The graph shows the average +/- the standard deviation of three independent experiments. 50nM wildtype Top2 (HFP185) or the ATPase dead E66Q mutant (HFP271) was incubated with or without 100μM salmon sperm DNA. (B) 15nM Uls1 (HFP350) and/or 50nM Top2 E66Q (HFP271) was incubated with or without 100μM salmon sperm DNA. Uls1 has weak DNA-stimulated ATPase activity which is not significantly further stimulated by Top2. Top2 E66Q was used to preferentially monitor the ATPase activity of Uls1.

**Supplementary figure S5. Top2 peak number increases in the presence of ACF.**

(A) Table showing the number of Top2 peaks associated with RNA Pol II genes, tRNA genes and replication origins (ARS) in WT (HFY250) or *uls1*Δ (HFY252) cells in the presence (ACF) or absence (YPD) of ACF. (B) ChIP qPCR (top panel) and ChIP-seq (bottom panel) at four different regions display the same overall trends +/- ACF. Top2 ChIP qPCR was performed on WT (HFY294) and *uls1*Δ cells (HFY297) where there is only one copy of *TOP2* and this is HA tagged. (C) Gene ontology analysis of regions that show a decrease in chromatin-bound Top2 after the addition of ACF in wildtype cells. Ribosomal protein genes are significantly enriched.

**Supplementary figure S6. Analysis of Top2 and Uls1 ChIP signal at repetitive loci**.

(A) Graph showing the number of Uls1 peaks associated with RNA Pol II genes, tRNA genes and replication origins (ARS) in WT (HF176) cells in the presence (ACF) or absence (YPD) of ACF. (B) Pairwise comparison of the average Uls1 ChIP enrichment using unfiltered reads across the genome and specifically within the rDNA locus, telomeric Y’ elements, tRNA genes and Ty retrotransposons +/- 100μM ACF. All pairwise comparisons (+/- ACF) with a Cohen’s *d* value > 0.2 are displayed. (C) Pairwise comparison of the average Top2 ChIP enrichment using unfiltered reads across the genome and specifically within the rDNA locus, telomeric Y’ elements, tRNA genes and Ty retrotransposons +/- 250μM ACF in either WT (HFY250) or *uls1*Δ (HFY252) cells. All pairwise comparisons (+/- ACF) with a Cohen’s *d* value > 0.2 are displayed.

